# Spatial and regional directory of tropical *Auricularia* mushrooms in Southwest, Nigeria

**DOI:** 10.1101/2021.05.07.443104

**Authors:** Victor S. Ekun, Clementina O. Adenipekun, Omena B. Ojuederie, Peter M. Etaware

## Abstract

Bioremediation of wastelands and dumpsites in Africa is fast declining due to reduced mushroom populations. In the past, the forest of Africa was teaming with mushrooms, but nowadays; mushrooms are severely exploited, resulting in gradual drift to extinction. Mushrooms have the tendency to degrade recalcitrant wastes and absorb heavy metals (Bio-accumulation). Unless concerted efforts are made to rejuvenate or rescue the surviving mushroom population, Africa will one day be overshadowed by wastes. The mushroom diversity in Southwest, Nigeria was determined by both morphological and molecular markers, 14 primers (OPB-11, OPB-12, OPB-15, OPB-20, OPB-21, OPH-3, OPH-5, OPH-10, OPH-15, OPT-1, OPT-5, OPT-7, OPT-10 and OPT-19) produced polymorphism with the test samples under electrophoresis gel (PCR and RAPD). Using standard morphological markers, *Auricularia auricula* was found to be evenly distributed across 8 locations in Ekiti and Osun, 6 locations in Ogun, 5 locations in Oyo and 4 locations in Lagos. There was none identified in Ondo. *Auricularia polytricha* was found in abundance in all the locations in Ondo. Lagos only had 3 out of its outline Stations graced with the presence of *A. polytricha*, whereas, Ogun, Ekiti, Osun and Oyo had no records of *A. polytricha*. From the genetic dissimilarity chart, 6 clusters of mushroom, sub-characterized into 3 distinct species (*Auricularia polytricha*, *A. auricula* and an unrelated *Auricularia* outlier species) and 5 cultivars were obtained in the region of Southwest, Nigeria. The population of all the *Auricularia* mushrooms currently present in Southwest, Nigeria was effectively captioned (Location, type and identity) by this research.

## 1.0 Introduction

Mushroom foraging was once considered as a prominent means of generating income for the locals of West Africa and it was also a pertinent raw material in folklore and traditional medicine (Guissou *et al*., 2008). In the past, West Africa was regarded as a major hotspot of mushroom diversity (Hawksworth, 2004), and as such, the implementation of mushrooms in the bioremediation of wastelands and dumpsites was considered as a lucrative and environmentally friendly approach, because mushrooms have the tendency to degrade ligno-cellulosic wastes within a short period of time and even possess the capacity to bio-accumulate and harness a vast array of heavy metals from the soil (Adenipekun *et al*., 2015). Mushrooms are majorly found in the wild (Osemwegie *et al*., 2014), or close to human settlements (Crous *et al*., 2006).

Mushrooms are currently threatened by extinction due to the massive destruction of their natural habitat (The wild) by natural disasters e.g. bush fire, earth quake, tornado etc., or by human intervention for their selfish interest e.g. construction of schools, hospitals, industries, houses, sport centres etc. (Gateri *et al*., 2004) and overgrazing by animals/humans. Mushroom domestication was the first line of action taken by ancient man to secure, safeguard, preserve and conserve mushroom species from extinction. One of the most economically important genera of the mushroom family is “*Auricularia* mushroom”, they are mostly edible mushroom of global repute with only about 17% global production, currently ranked 3^rd^ as the most cultivated mushroom genus after *Lentinula* (22%) and *Pleurotus* (19%) (Royse *et al*., 2017; Bandara *et al*., 2019).

The genus *Auricularia* belongs to the family *Auriculariaceae*, class *Agaricomycetes*, phylum *Basidiomycota* and kingdom *Fungi* (Moore *et al*., 2001; Moore 2013). Several species exist within the fungi order “*Auriculariales*” that are useful as both edible and medicinal mushrooms (Chang and Hayes, 2013). *Auricularia* mushrooms are widespread throughout the temperate and sub-tropical zones of the world, and can be found across Europe, North America, Asia, and Australia (Conte and Laessoe, 2008). It has been reported that there are only 15-20 species of *Auricularia* worldwide with 8 species identified in China (Chang and Miles, 2004). Among these species, *A. auricula and A. polytricha* are the most popular and the most cultivated around the world (Chang and Miles, 2004). *Auricularia auricula* commonly known as wood ear mushrooms is native to Kenya and occurs in Kakamega forest in Western Kenya. In other parts of Africa, the wood ear mushrooms have been reported in Nigeria where it is being conserved through cultivation on palm substrates (Osemwegie and Okhuoya, 2009). In Kenya, the wood ears have not been previously cultivated because they are protected by wildlife conservation laws.

According to Osemwegie *et al*. (2014), proper inventory of wild or domesticated edible mushrooms with high medicinal values, sold in local markets is required for the development of a mushroom genetic resource or germplasm, a database for differentiating between toxic and edible mushrooms, and the cultivation of species yet uncultivated. Above all, the safety of life and a decline in the cases of “Mycotoxicosis” resulting in human and wildlife casualty is of paramount global interest. The dearth of information regarding the domestication and cultivation of mushrooms (Mushroom technology) became the major cause for dependence on mushroom hunting/foraging, practiced by the indigenous people of Africa. The common way to identify different *Auricularia* species was based on morphological characters such as size, shape and colour of the fruiting body etc. Musngi *et al*. (2004) effectively classified various strains of *Auricularia* spp in the Philippines by simply using phenotypic characters.

The use of morphological markers only for characterization of *Auricularia* species found in Southwest, Nigeria is largely unreliable, misleading and ineffective because it has several limitations which abound majorly due to the adverse effects and intricate influence of environmental factors (Etaware *et al*., 2020) on the phenotype or physical appearance of similar mushrooms species grown under different environmental conditions. Therefore, the use of molecular markers (PCR and RAPD) to determine the genetic diversity and variation among large genomic entity of *Auricularia* mushrooms with similar gene pool is an added advantage, a more reliable and valuable tool that can detect the slightest trace of genetic variability, which is the basis for characterization and classification of these species into a more organized taxa (Al-Gabbiesh *et al*., 2006).

Finally, a comprehensive knowledge of the varietal differences that exist within the genus, species and sub-species of the *Auricularia* mushroom can be exploited by the use of molecular markers, which will inferably serve as sources of cell lines for researchers and in-breeding programs within Africa and around the world (Pei-Sheng and Chang, 2004). Inferably without an aorta of doubt, one can surmise that the inability to effectively distinguish between poisonous and edible mushroom may also have accounted for the visible underdevelopment in global mushroom cultivation which has undermined the commercial scale production of edible and medicinal mushrooms for decades unending leading to low priority export or impact on foreign exchange earnings.

## 2.0 Materials and Methods

### 2.1 Spatial grouping (Stereotype and Catalogue) of *Auricularia* sp

A total of fifty four (54) sample stations was setup across Southwest, Nigeria for the sole purpose of efficiency and accuracy in the catalogue of wild *Auricularia* mushrooms in the tropical, sub-tropical, and rain forest region of Southwest, Nigeria (See Table 1–6 for comprehensive details). Field assessment was conducted regularly between September 2011 and July 2012. The geographical view of Southwest, Nigeria was described in Fig 1. The coordinates and geographical identity of each sample station was referenced in Table 1–6. Prospective *Auricularia* samples were identified at the Department of Botany, University of Ibadan, Ibadan, Oyo State, Nigeria, based on the recommended phenotypic behaviours (colour, shape, texture, and fruit body) given by Musngi *et al*. (2004).

**Fig 1.**
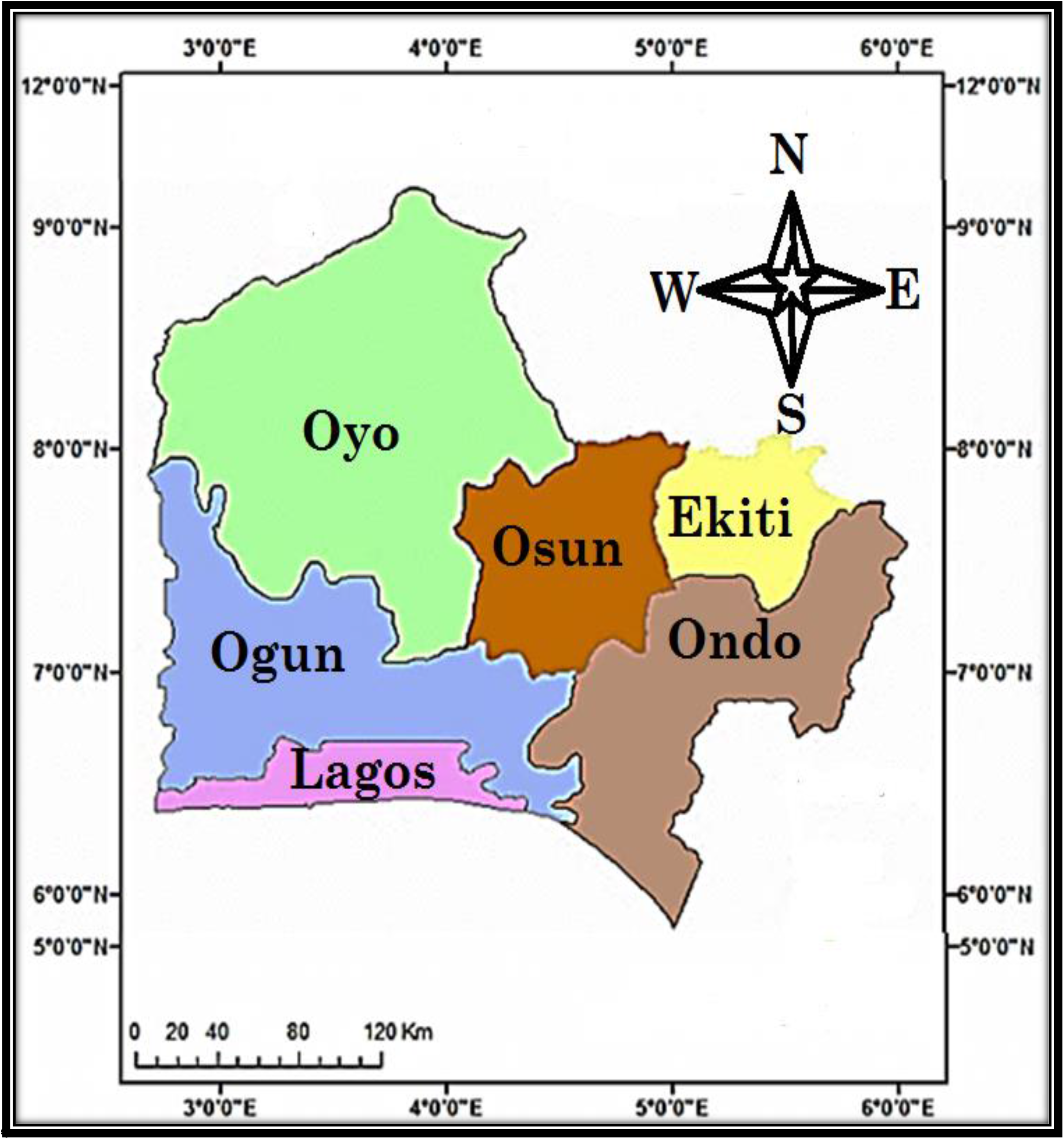
The geographical positioning of the States that comprises Southwestern Nigeria

**Table 1:** The coordinates and location of sample stations in Ogun State, Southwest-Nigeria

| S/N | Stations | Code | Local Govt. Area | Town | State | Latitude | Longitude |
| --- | --- | --- | --- | --- | --- | --- | --- |
| 1 | 01 | OG1 | Abeokuta North | Abeokuta | Ogun | 7.1475°N | 3.3619°E |
| 2 | 02 | OG2 | Ewekoro | Itori | Ogun | 6.9530°N | 3.2181°E |
| 3 | 03 | OG3 | Ifo | Ifo | Ogun | 6.8192°N | 3.1930°E |
| 4 | 04 | OG4 | Ijebu Ode | Ijebu Ode | Ogun | 6.8300°N | 3.9165°E |
| 5 | 05 | OG5 | Ikenne | Ikenne | Ogun | 6.8717°N | 3.7105°E |
| 6 | 06 | OG6 | Shagamu | Shagamu | Ogun | 6.8322°N | 3.6319°E |
| 7 | 07 | OG7 | Odeda | Odeda | Ogun | 7.2328°N | 3.5281°E |
| 8 | 08 | OG8 | Odogbolu | Odogbolu | Ogun | 6.8365°N | 3.7689°E |

State Code: OG→Ogun → No *Auricularia* specimen found in that region

**Table 2:** The coordinates and location of sample stations in Lagos State, Southwest-Nigeria

| S/N | Stations | Code | Local Govt. Area | Town | State | Latitude | Longitude |
| --- | --- | --- | --- | --- | --- | --- | --- |
| 1 | 09 | LA1 | Agege | Ikeja | Lagos | 6.6180°N | 3.3209°E |
| 2 | 10 | LA2 | Ojo | Ojo | Lagos | 6.4579°N | 3.1580°E |
| 3 | 11 | LA3 | Apapa | Ikeja | Lagos | 6.4553°N | 3.3641°E |
| 4 | 12 | LA4 | Badagry | Badagry | Lagos | 6.4316°N | 2.8876°E |
| 5 | 13 | LA5 | Epe | Epe | Lagos | 6.6055°N | 3.9470°E |
| 6 | 14 | LA6 | Shomolu | Shomolu | Lagos | 6.5392°N | 3.3842°E |
| 7 | 15 | LA7 | Ikorodu | Ikorodu | Lagos | 6.6194°N | 3.5105°E |
| 8 | 16 | LA8 | Mushin | Ikeja | Lagos | 6.5273°N | 3.3414°E |

State Code: LA→Lagos → No *Auricularia* specimen found in that region

**Table 3:** The coordinates and location of sample stations in Oyo State, Southwest-Nigeria

| S/N | Stations | Code | Local Govt. Area | Town | State | Latitude | Longitude |
| --- | --- | --- | --- | --- | --- | --- | --- |
| 1 | 17 | OY1 | Akinyele | Moniya | Oyo | 7.5249°N | 3.9152°E |
| 2 | 18 | OY2 | Egbeda | Egbeda | Oyo | 7.3796°N | 3.9675°E |
| 3 | 19 | OY3 | Ido | Ido | Oyo | 7.5077°N | 3.7194°E |
| 4 | 20 | OY4 | Iseyin | Iseyin | Oyo | 7.9765°N | 3.5914°E |
| 5 | 21 | OY5 | Ogbomosho North | Ogbomosho | Oyo | 8.1227°N | 4.2436°E |
| 6 | 22 | OY6 | Oluyole | Idi Ayunre | Oyo | 7.2247°N | 3.8732°E |
| 7 | 23 | OY7 | Oyo | Oyo | Oyo | 7.8430°N | 3.9368°E |
| 8 | 24 | OY8 | Olorunsogo | Igbeti | Oyo | 8.7699°N | 4.1104°E |
| 9 | 52 | OY9 | Akinyele | Ojo | Oyo | 7.5503°N | 3.9470°E |
| 10 | 53 | OY10 | Ibadan North | Bodija | Oyo | 7.4351°N | 3.9143°E |

State Code: OY→Oyo → No *Auricularia* specimen found in that region

**Table 4:** The coordinates and location of sample stations in Ekiti State, Southwest-Nigeria

| S/N | Stations | Code | Local Govt. Area | Town | State | Latitude | Longitude |
| --- | --- | --- | --- | --- | --- | --- | --- |
| 1 | 25 | EK1 | Ado Ekiti | Ado Ekiti | Ekiti | 7.6124°N | 5.2371°E |
| 2 | 26 | EK2 | Ilejemeje | Iye | Ekiti | 7.9591°N | 5.2371°E |
| 3 | 27 | EK3 | Ikole | Ikole | Ekiti | 7.7983°N | 5.5145°E |
| 4 | 28 | EK4 | Oye | Oye | Ekiti | 7.7979°N | 5.3286°E |
| 5 | 29 | EK5 | Irepodun | Igede | Ekiti | 7.7313°N | 5.2476°E |
| 6 | 30 | EK6 | Ikere | Ikere | Ekiti | 7.4991°N | 5.2319°E |
| 7 | 31 | EK7 | Ijero | Ijero Ekiti | Ekiti | 7.8120°N | 5.0677°E |
| 8 | 32 | EK8 | Emure | Emure Ekiti | Ekiti | 7.4317°N | 5.4621°E |

State Code: EK→Ekiti → No *Auricularia* specimen found in that region

**Table 5:** The coordinates and location of sample stations in Ondo State, Southwest-Nigeria

| S/N | Stations | Code | Local Govt. Area | Town | State | Latitude | Longitude |
| --- | --- | --- | --- | --- | --- | --- | --- |
| 1 | 33 | OD1 | Idanre | Idanre | Ondo | 7.0914°N | 5.1484°E |
| 2 | 34 | OD2 | Ilaje | Igbokoda | Ondo | 6.2585°N | 4.7692°E |
| 3 | 35 | OD3 | Ile Oluji | Ile Oluji | Ondo | 7.2825°N | 4.8521°E |
| 4 | 36 | OD4 | Odigbo | Ore | Ondo | 6.7519°N | 4.8780°E |
| 5 | 37 | OD5 | Okitipupa | Okitipupa | Ondo | 6.5025°N | 4.7795°E |
| 6 | 38 | OD6 | Ose | Ifon | Ondo | 6.9235°N | 5.7774°E |
| 7 | 39 | OD7 | Owo | Owo | Ondo | 7.1989°N | 5.5932°E |
| 8 | 40 | OD8 | Ifedore | Igbara Oke | Ondo | 7.3877°N | 5.0807°E |
| 9 | 54 | OD9 | Akure South | Akure | Ondo | 7.2571°N | 5.2058°E |

State Code: OD→Ondo → No *Auricularia* specimen found in that region

**Table 6:**
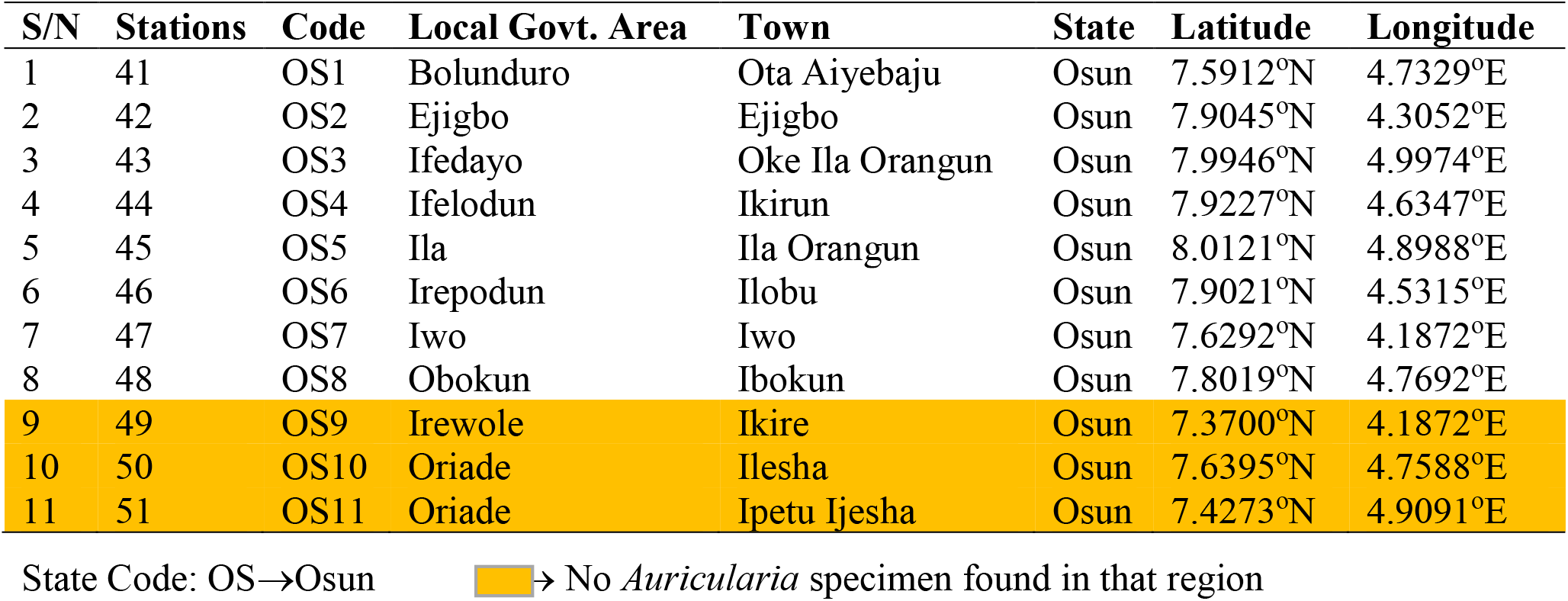
The coordinates and location of sample stations in Osun State, Southwest-Nigeria

### 2.2 Morphological characterization of prospective *Auricularia* sp

The basidiocarps were rehydrated by soaking in water for 10 minutes before characterization. Qualitative characters such as colour, shape, and presence of hymenia was evaluated by physical observation while texture was determined by touching the back and top surfaces (Onyango *et al*., 2011). For microscopic characters, free hand transverse sections of approximately 0.1 mm thick were made from rehydrated basidiocarps with the aid of a sharp surgical blade. The sections were immersed in a diluted solution of methylene blue stain and left for 10 minutes. Thin sections were selected and placed on glass slides, fitted with cover slips and the anatomy of each basidiocarp was studied. The characterization of *Auricularia* spp based on morphological markers (Colour, Shape, and Texture etc. of both the mycelia and entire mushroom body) was described in Table 7.

**Table 7:** Classification of Auricularia species using morphological markers

| Case | External Mushroom Features |  |  | Mycelia Features |  | Inference |  |
| --- | --- | --- | --- | --- | --- | --- | --- |
|  | Colour | Shape | Texture | Colour | Type | Genus | Species |
| 1 | Dark brown | Discoid | Gelatinous | White | Cottony | *Auricularia* | Unknown |
| 2 | Yellow brown | Auriform | Leathery | Off white | Cottony | *Auricularia* | *auricula* |
| 3 | Brown | Flattened | Rubbery | Off white | Scanty | *Auricularia* | *Unknown* |
| 4 | Dark brown | Discoid | Gelatinous | White | Cottony | *Auricularia* | *polytricha* |

A modified protocol adapted from Onyango *et al.* (2011)

### 2.3 Molecular characterization of prospective *Auricularia* sp

#### 2.3.1 Tissue preparation and DNA Extraction

The modified DNA extraction protocol of Chen *et al*. (2010), involving the use of Cetyltrimethylammonium Bromide (CTAB), was used for DNA isolation. Tissues from the pileus of each mushroom specimens was aseptically detached using a sterile scalpel and 200mg was weighed out prior to DNA extraction. The tissues was pulverized with 800ml of CTAB buffer (20 mM EDTA, 1.4 mM NaCl, 100 mM Tris-HCl pH 8.0, SDS (1.25%, 2% CTAB and 0.2% β-mercaptoethanol (v/v)), incubated at 65°C for 15 min using water bath with intermittent homogenization, allowed to cool for approximately 1 min before adding equal volume of phenol, chloroform and iso-amyl alcohol at the ratio of 25:24:1.

The mixture was further centrifuged at 12,000 revolutions per minute (rpm) for 15 minutes; the supernatant was transferred to clean sterile tubes without unsettling the pellets. About 400 μl of ice-cold isopropanol was added to the supernatant and mixed by inverting the tubes 2-5 times to precipitate the DNA and subsequently kept at −80°C for 1hr. The DNA sediment was pelleted by centrifugation at 12,000 rpm for 10 min and the dried DNA pellets obtained were re-suspended in 100 μl of Grand Island Biological Company (GIBCO) water (Invitrogen, Carlsbad, CA, USA) and 2 μl of 10 mg/ml RNase (Qiagen Valencia, CA, USA) was added to each of the samples and kept at 4°C for 30 minutes to remove fragments of RNA strands.

#### 2.3.2 DNA Sequencing

The extracted DNA fragments from each Auricularia mushroom specimen was sequenced using the high-throughput Sequencing (HTS) or Next Generation Sequencing (NGS) Technique. A total of 2.5μl of the stock DNA samples were loaded on 1.5% agarose gel for electrophoresis and visualized under UV light (Model-2, Upland, CA, USA) to check the quality of the extracted DNA samples. Following the high level of concentration of the extracted DNA samples, dilution of each DNA sample was uniformly made to 100ng/uL DNA.

#### 2.3.3 DNA Purification and Quantification

The sequenced DNA fragments were quantified using Nano-Drop spectrophotometer (ND-1000). About 2μl of the extracted DNA sample was used to obtain a unique ration of 1.8:2.0 at OD 260/280 absorbance level and concentration through which dilution samples were prepared for polymerase chain reaction (PCR).

*Mathematically*,

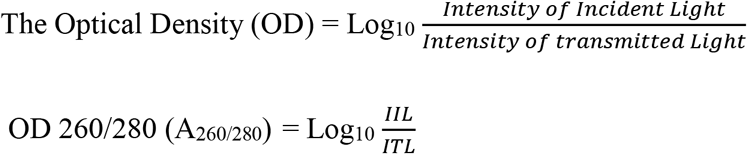

#### 2.3.4 DNA/RNA primer selection and buffer preparation

A total of twenty five (25) primers were subjected to screening for polymorphism with the prospective *Auricularia* species (i.e. OPB-1, OPB-2, OPB-3, OPB-4, OPB-5, OPB-6, OPB-7, OPB-8, OPB-9, OPB-10, OPB-11, OPB-12, OPB-15, OPB-20, OPB-21, OPH-3, OPH-5, OPH-10, OPH-15, OPT-1, OPT-5, OPT-7, OPT-10, OPT-19 and OPD-18) out of which fourteen (14) primers were polymorphic (OPB-11, OPB-12, OPB-15, OPB-20, OPB-21, OPH-3, OPH-5, OPH-10, OPH-15, OPT-1, OPT-5, OPT-7, OPT-10 and OPT-19) as shown in Table 8. The fourteen (14) arbitrary RAPD decamer primers (Table 8) obtained from Operon Technology (Alameda, CA, USA) were used for PCR amplification.

**Table 8:** Primers used for DNA amplification during molecular analysis

| S/N | RAPD primer | DNA/RNA Primer Sequence | Melting Point (Tm°C) |
| --- | --- | --- | --- |
| 1 | OPB-11 | 5<sup>1</sup>GTAGACCCGT 3<sup>1</sup> | 34 |
| 2 | OPB-12 | 5<sup>1</sup>CGTTGACGCA 3<sup>1</sup> | 34 |
| 3 | OPB-15 | 5<sup>1</sup>GGAGGGTGTT 3<sup>1</sup> | 32 |
| 4 | OPB-20 | 5<sup>1</sup>GGACCCTTAC 3<sup>1</sup> | 34 |
| 5 | OPB-21 | 5<sup>1</sup>CGACCCTTAC 3<sup>1</sup> | 34 |
| 6 | OPH-3 | 5<sup>1</sup>AGACGTCCAC 3<sup>1</sup> | 34 |
| 7 | OPH-5 | 5<sup>1</sup>AGTCGTCCCC 3<sup>1</sup> | 32 |
| 8 | OPH-10 | 5<sup>1</sup>CCTACGTCAG 3<sup>1</sup> | 32 |
| 9 | OPH-15 | 5<sup>1</sup>GCTTCGTCAG 3<sup>1</sup> | 34 |
| 10 | OPT-1 | 5<sup>1</sup>GGGCCACTCA 3<sup>1</sup> | 34 |
| 11 | OPT-5 | 5<sup>1</sup>GGGTTTGGCA 3<sup>1</sup> | 32 |
| 12 | OPT-7 | 5<sup>1</sup>GGCAGGCTGT 3<sup>1</sup> | 34 |
| 13 | OPT-10 | 5<sup>1</sup>CCTTCGGAAG 3<sup>1</sup> | 32 |
| 14 | OPT-19 | 5<sup>1</sup>GATGCCAGAC 3<sup>1</sup> | 32 |

Operon Technology, Alameda, California, USA

#### 2.3.5 PCR Analysis (DNA Amplification and Fingerprinting)

The required capacity for PCR amplification was 25μl i.e. 2.0μl of 100ng DNA, 2.5μl of 10 x Buffer (Bioline), 1.25μl of 50mM MgCl_2_ (Bioline), 2.0μl of 2.5mM dNTPs (Bioline), and 0.2μl 500U *Taq* DNA polymerase (Bioline), 1.0μl DMSO (dimethyl sulfoxide), 1.0μl of 10uM of each primer and 16.05μl of 500ml DEPC-treated water (Invitrogen Corporation). PCR amplifications were performed using Applied Bio-systems thermo-cycler with a cycling profile of an initial step of 94°C for 2 minutes, 40 cycles of 94°C for 20 s, 72°C for 1min, and 54°C for 2 mins., and a 5-min final extension at 72°C.

#### 2.3.6 RAPD profiling using electrophoresis gel

Amplified fragments were separated by electrophoresis on 1.5% (w/v) agarose (Sigma Aldrich, USA) gels with 1X TBE (Tris-Boric acid-EDTA) buffer and stained with ethidium bromide (0.5mg/ml). The molecular fragments were estimated using 100-bp step DNA marker (Bio-labs, New England).

### 2.4 Statistical Analysis

Data matrix generated from the RAPD sequence for fragments of similar molecular weight from each individual mushroom specimens were scored as present (1) or absent (0). The data obtained from scoring the RAPD bands were used to determine the genetic dissimilarity matrix using Jaccard’s similarity coefficient (Jaccard 1908 Standard Protocol). Phylogenetic relations were determined by cluster analysis using UGPMA (un-weighted pair-group method with arithmetic averages) aided by the NTSYS-pc software version 2.02 (Rohlf 1998 Preferred Protocol). Phylogenetic characterization into multivariate groups was done using principal component analysis (PCA) with Darwin software version 5.0.0.157 while polymorphic information content (PIC) was calculated using the method of Botstein *et al*. (1980). The data obtained were analyzed using a one way analysis of variance (ANOVA) aided by SPSS v20. Significantly different means were separated using Tukey test at P<0.05

## 3.0 Results

### 3.1 Geo-mapping of *Auricularia* sp. in Southwest, Nigeria

A total of 31 samples of *Auricularia auricula* were identified and geo-tagged at several strategic locations within Southwest, Nigeria (See Table 9–14 for more details). *Auricularia auricula* was evenly distributed across 8 sample Stations in Ekiti (Table 12) and Osun (Table 14) States, 6 locations in Ogun State (Table 9), 5 Stations in Oyo State (Table 11) and 4 locations in Lagos State (Table 10). There was none identified in Ondo State (Table 13) as at the time of filing this reports. *Auricularia polytricha* was found in abundance in Ondo State i.e. it was evenly distributed around 8 strategic locations within the State (Table 13). Lagos State only had 3 out of its outline Stations graced with the presence of *A. polytricha* (Table 10). Ogun, Ekiti, Osun and Oyo States had no records of *A. polytricha* i.e. the mushroom was not found within their forest domain prior to the compilation of this report. About 5 species from the genus *Auricularia* found in Ogun (Table 9), Lagos (Table 10) and Oyo (Table 11) States were not identified to their species level due to discrepancy in their morphological status.

**Table 9:**
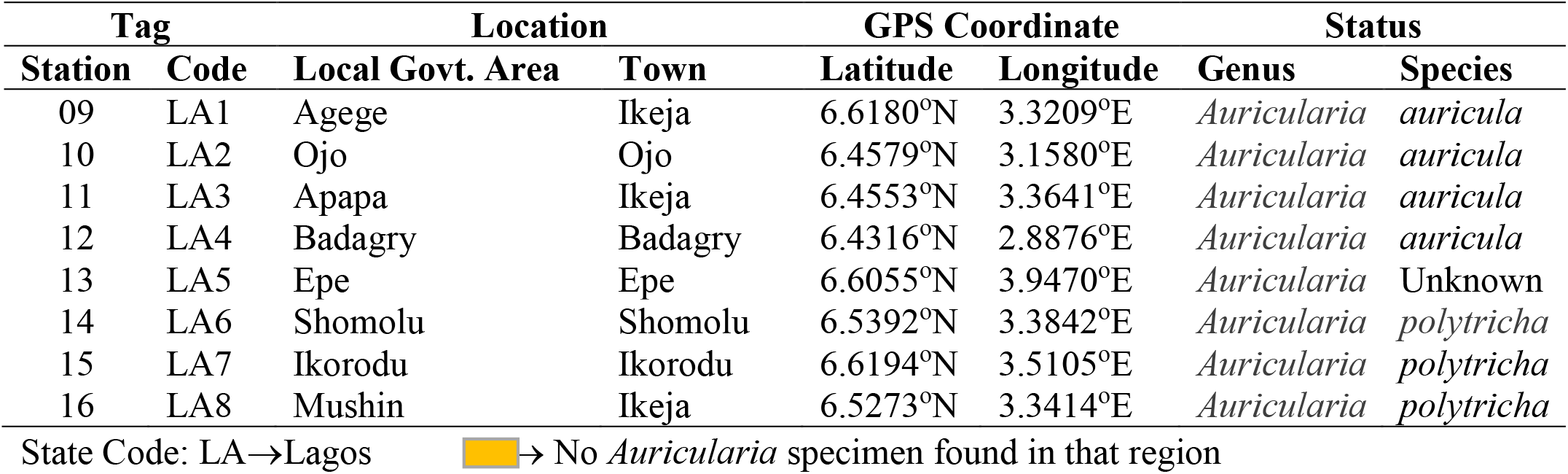
Geo-mapping of *Auricularia* species in Ogun, Nigeria based on morphological characters

**Table 10:**
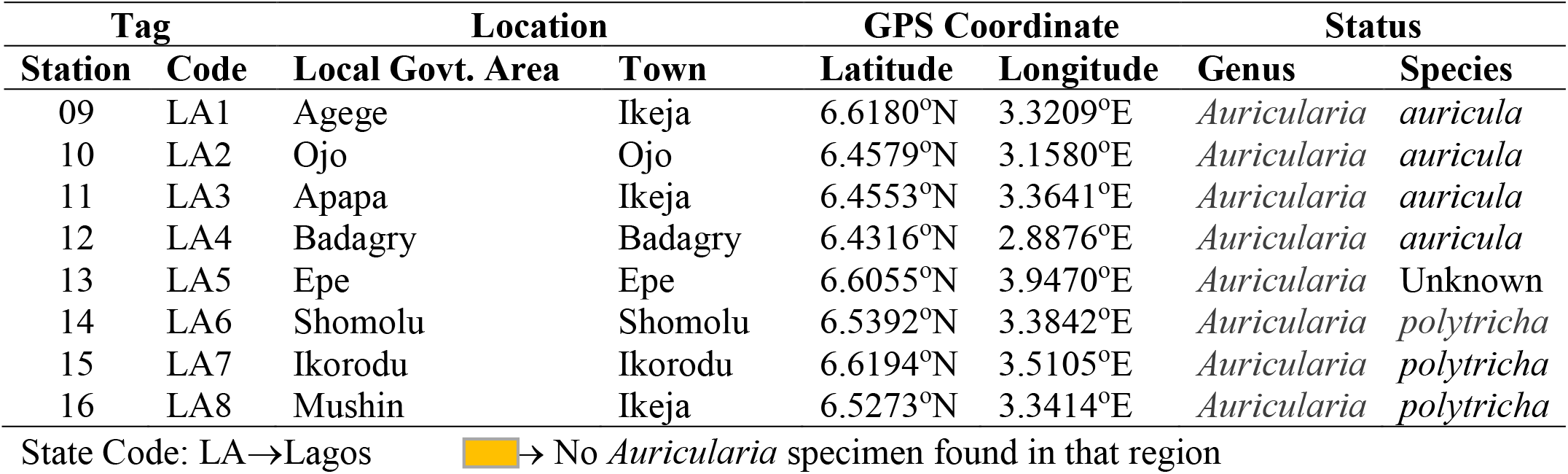
Geo-mapping of *Auricularia* species in Lagos, Nigeria based on morphological characters

**Table 11:** Geo-mapping of *Auricularia* species in Oyo, Nigeria based on morphological characters

| Tag |  | Location |  | GPS Coordinate |  | Status |  |
| --- | --- | --- | --- | --- | --- | --- | --- |
| Station | Code | Local Govt. Area | Town | Latitude | Longitude | Genus | Species |
| 17 | OY1 | Akinyele | Moniya | 7.5249°N | 3.9152°E | *Auricularia* | *polytricha* |
| 18 | OY2 | Egbeda | Egbeda | 7.3796°N | 3.9675°E | *Auricularia* | *auricula* |
| 19 | OY3 | Ido | Ido | 7.5077°N | 3.7194°E | *Auricularia* | Unknown |
| 20 | OY4 | Iseyin | Iseyin | 7.9765°N | 3.5914°E | *Auricularia* | Unknown |
| 21 | OY5 | Ogbomosho North | Ogbomosho | 8.1227°N | 4.2436°E | *Auricularia* | *auricula* |
| 22 | OY6 | Oluyole | Idi Ayunre | 7.2247°N | 3.8732°E | *Auricularia* | *auricula* |
| 23 | OY7 | Oyo | Oyo | 7.8430°N | 3.9368°E | *Auricularia* | *auricula* |
| 24 | OY8 | Olorunsogo | Igbeti | 8.7699°N | 4.1104°E | *Auricularia* | *auricula* |
| 52 | OY9 | Akinyele | Ojo | 7.5503°N | 3.9470°E | None | None |
| 53 | OY10 | Ibadan North | Bodija | 7.4351°N | 3.9143°E | None | None |

State Code: OY→Oyo → No *Auricularia* specimen found in that region

**Table 12:** Geo-mapping of *Auricularia* species in Ekiti, Nigeria based on morphological characters

| Tag |  | Location |  | GPS Coordinate |  | Status |  |
| --- | --- | --- | --- | --- | --- | --- | --- |
| Station | Code | Local Govt. Area | Town | Latitude | Longitude | Genus | Species |
| 25 | EK1 | Ado Ekiti | Ado Ekiti | 7.6124°N | 5.2371°E | *Auricularia* | *auricula* |
| 26 | EK2 | Ilejemeje | Iye | 7.9591°N | 5.2371°E | *Auricularia* | *auricula* |
| 27 | EK3 | Ikole | Ikole | 7.7983°N | 5.5145°E | *Auricularia* | *auricula* |
| 28 | EK4 | Oye | Oye | 7.7979°N | 5.3286°E | *Auricularia* | *auricula* |
| 29 | EK5 | Irepodun | Igede | 7.7313°N | 5.2476°E | *Auricularia* | *auricula* |
| 30 | EK6 | Ikere | Ikere | 7.4991°N | 5.2319°E | *Auricularia* | *auricula* |
| 31 | EK7 | Ijero | Ijero Ekiti | 7.8120°N | 5.0677°E | *Auricularia* | *auricula* |
| 32 | EK8 | Emure | Emure Ekiti | 7.4317°N | 5.4621°E | *Auricularia* | *auricula* |

State Code: EK→Ekiti → No *Auricularia* specimen found in that region

**Table 13:**
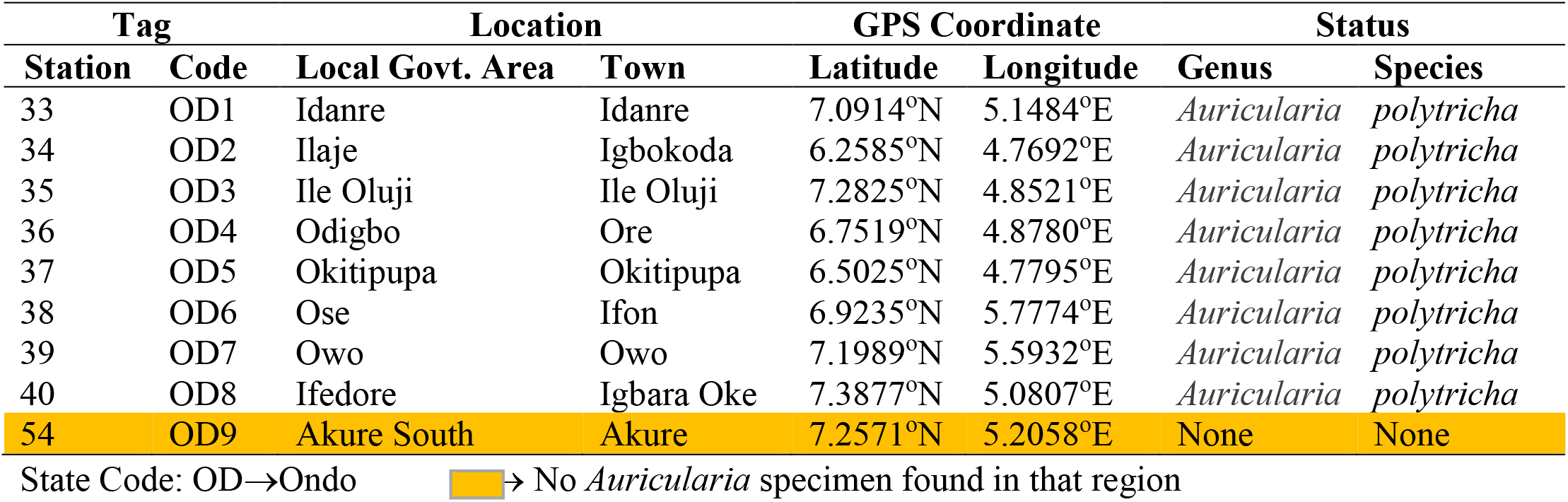
Geo-mapping of *Auricularia* species in Ondo, Nigeria based on morphological characters

| Tag |  | Location |  | GPS Coordinate |  | Status |  |
| --- | --- | --- | --- | --- | --- | --- | --- |
| Station | Code | Local Govt. Area | Town | Latitude | Longitude | Genus | Species |
| 33 | OD1 | Idanre | Idanre | 7.0914°N | 5.1484°E | *Auricularia* | *polytricha* |
| 34 | OD2 | Ilaje | Igbokoda | 6.2585°N | 4.7692°E | *Auricularia* | *polytricha* |
| 35 | OD3 | Ile Oluji | Ile Oluji | 7.2825°N | 4.8521°E | *Auricularia* | *polytricha* |
| 36 | OD4 | Odigbo | Ore | 6.7519°N | 4.8780°E | *Auricularia* | *polytricha* |
| 37 | OD5 | Okitipupa | Okitipupa | 6.5025°N | 4.7795°E | *Auricularia* | *polytricha* |
| 38 | OD6 | Ose | Ifon | 6.9235°N | 5.7774°E | *Auricularia* | *polytricha* |
| 39 | OD7 | Owo | Owo | 7.1989°N | 5.5932°E | *Auricularia* | *polytricha* |
| 40 | OD8 | Ifedore | Igbara Oke | 7.3877°N | 5.0807°E | *Auricularia* | *polytricha* |
| 54 | OD9 | Akure South | Akure | 7.2571°N | 5.2058°E | None | None |

State Code: OD→Ondo → No *Auricularia* specimen found in that region

**Table 14:**
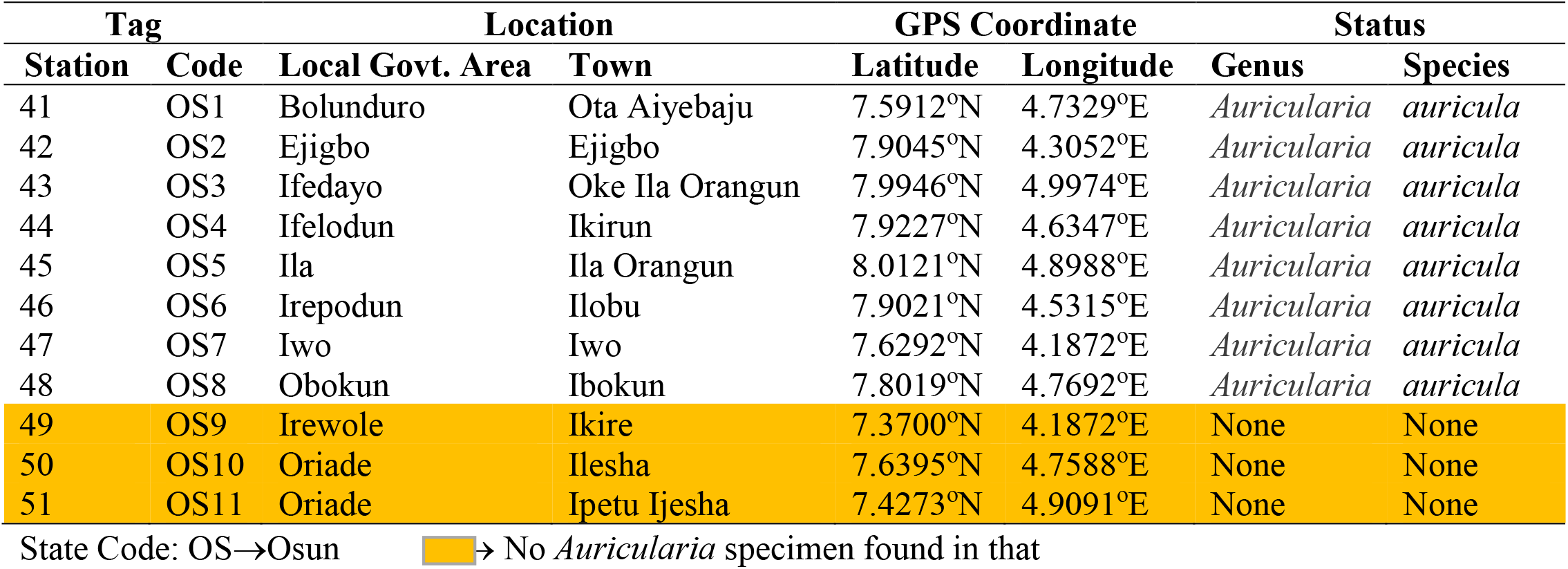
Geo-mapping of *Auricularia* species in Osun, Nigeria based on morphological characters

### 3.2 Molecular characterization of Prospective *Auricularia* sp

#### 3.2.1 DNA Purification (Quality) and Quantification

The prospective *Auricularia* specimens marked out from forty eight (48) locations within Southwest, Nigeria were further subjected to molecular test in order to ascertain and fully establish the genomic differences that exist among the mushroom specimens based on the influence of the environment and geographical boundaries, and further enhance the characterization made in this research based on morphological markers (Table 9–14). The first step was to extract and sequence their DNA materials. The extracted DNA from each prospective *Auricularia* Mushroom was tested for impurities; the purity of the extracted DNA samples was determined by UV light Absorbance at 260/280nm ratio using a spectrophotometer prior to PCR and RAPD analysis (See Table 15–20 for more details).

**Table 15:**
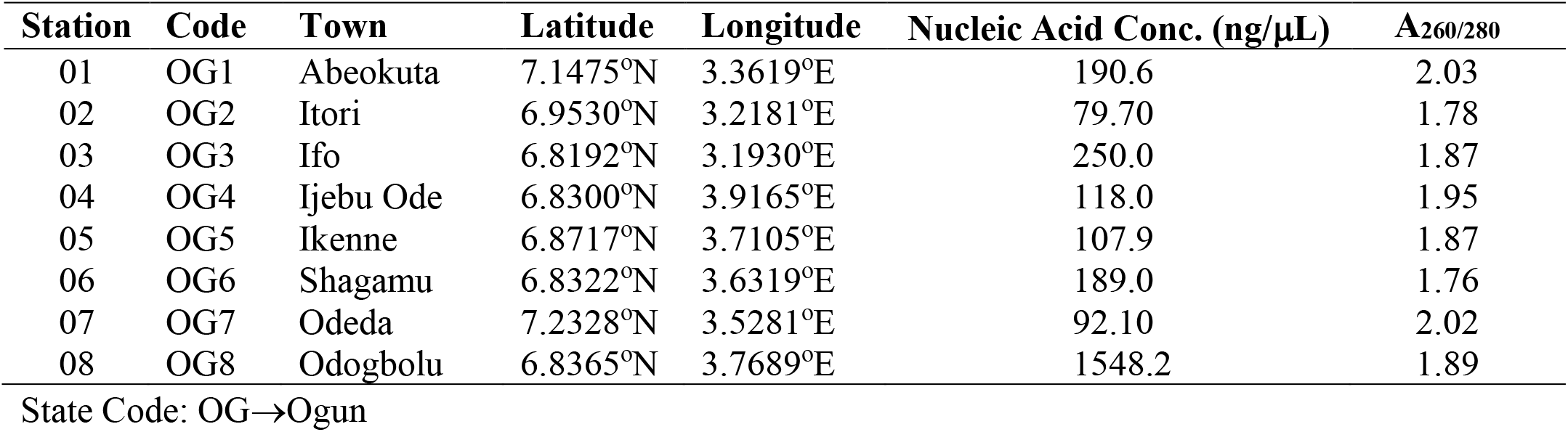
Qualitative assessment of nucleic acid extracted from *Auricularia* species in Ogun State

**Table 16:** Qualitative description of nucleic acid extracted from *Auricularia* species in Lagos State

| Station | Code | Town | Latitude | Longitude | Nucleic Acid Conc. (ng/μL) | A<sub>260/280</sub> |
| --- | --- | --- | --- | --- | --- | --- |
| 09 | LA1 | Ikeja | 6.6180°N | 3.3209°E | 79.00 | 2.08 |
| 10 | LA2 | Ojo | 6.4579°N | 3.1580°E | 282.3 | 2.14 |
| 11 | LA3 | Ikeja | 6.4553°N | 3.3641°E | 309.1 | 2.12 |
| 12 | LA4 | Badagry | 6.4316°N | 2.8876°E | 137.8 | 2.13 |
| 13 | LA5 | Epe | 6.6055°N | 3.9470°E | 890.0 | 1.83 |
| 14 | LA6 | Shomolu | 6.5392°N | 3.3842°E | 96.90 | 2.11 |
| 15 | LA7 | Ikorodu | 6.6194°N | 3.5105°E | 94.00 | 2.11 |
| 16 | LA8 | Ikeja | 6.5273°N | 3.3414°E | 187.7 | 2.03 |

State Code: LA→Lagos

**Table 17:** Qualitative description of nucleic acid extracted from *Auricularia* species in Oyo State

| Station | Code | Town | Latitude | Longitude | Nucleic Acid Conc. (ng/μL) | A<sub>260/280</sub> |
| --- | --- | --- | --- | --- | --- | --- |
| 17 | OY1 | Moniya | 7.5249°N | 3.9152°E | 110.6 | 2.06 |
| 18 | OY2 | Egbeda | 7.3796°N | 3.9675°E | 1507.5 | 1.92 |
| 19 | OY3 | Ido | 7.5077°N | 3.7194°E | 96.70 | 2.07 |
| 20 | OY4 | Iseyin | 7.9765°N | 3.5914°E | 1118 | 1.96 |
| 21 | OY5 | Ogbomosho | 8.1227°N | 4.2436°E | 543.5 | 1.99 |
| 22 | OY6 | Idi Ayunre | 7.2247°N | 3.8732°E | 193.8 | 2.10 |
| 23 | OY7 | Oyo | 7.8430°N | 3.9368°E | 490.7 | 2.01 |
| 24 | OY8 | Igbeti | 8.7699°N | 4.1104°E | 239.3 | 2.10 |

State Code: OY→Oyo

**Table 18:**
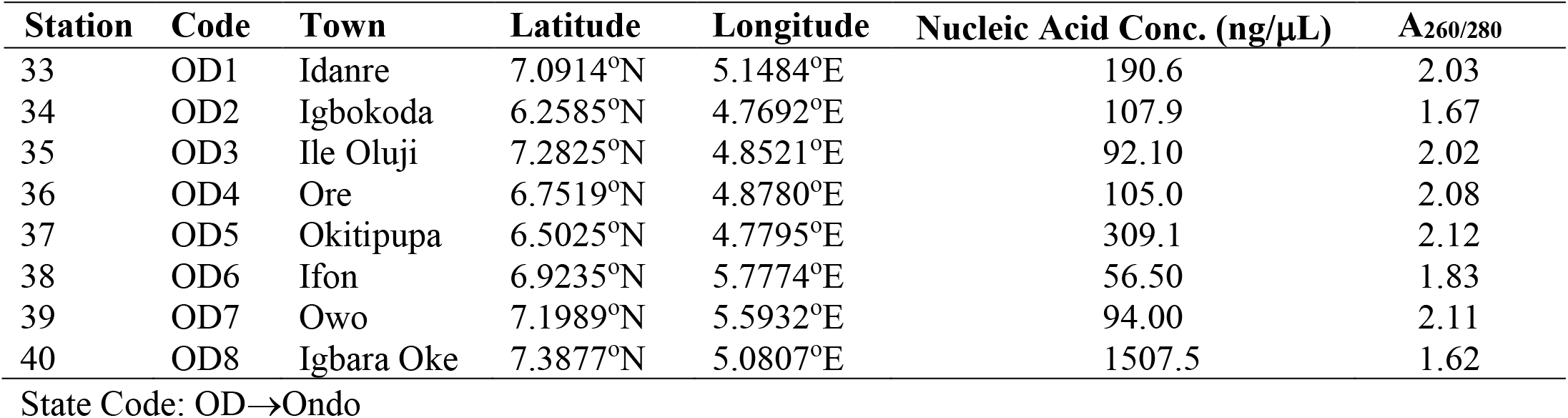
Qualitative description of nucleic acid extracted from *Auricularia* species in Ekiti State

**Table 19:**
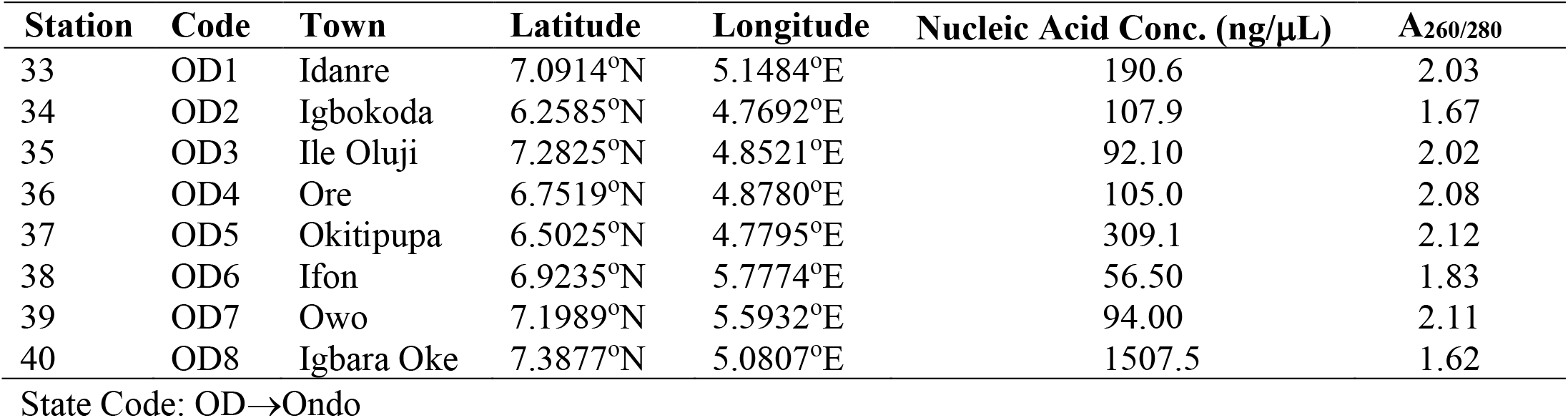
Qualitative description of nucleic acid extracted from *Auricularia* species in Ondo State

**Table 20:**
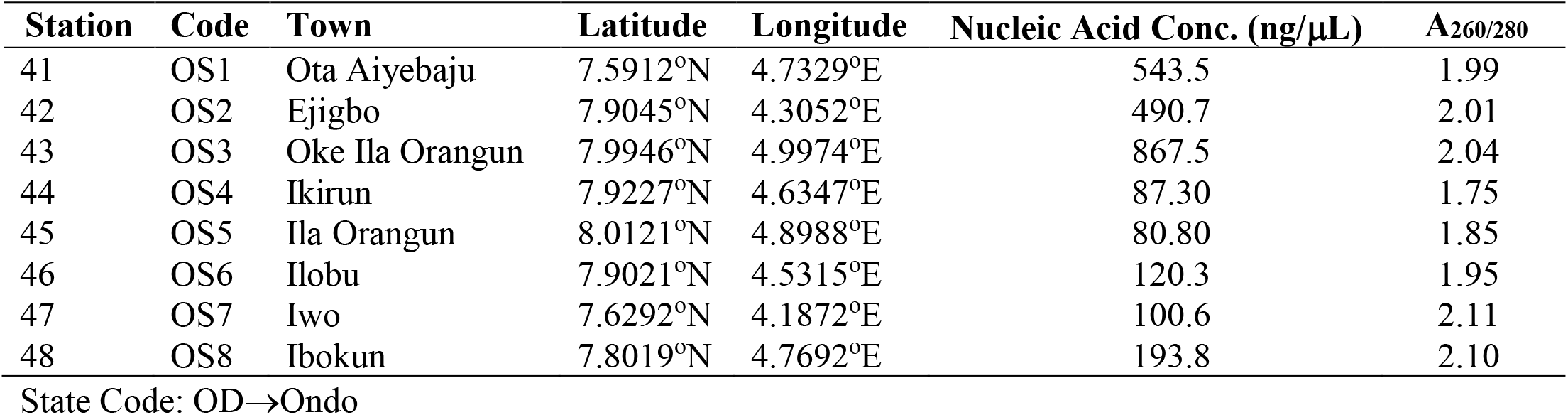
Qualitative description of nucleic acid extracted from *Auricularia* species in Osun State

| Station | Code | Town | Latitude | Longitude | Nucleic Acid Conc. (ng/μL) | A<sub>260/280</sub> |
| --- | --- | --- | --- | --- | --- | --- |
| 41 | OS1 | Ota Aiyebaju | 7.5912°N | 4.7329°E | 543.5 | 1.99 |
| 42 | OS2 | Ejigbo | 7.9045°N | 4.3052°E | 490.7 | 2.01 |
| 43 | OS3 | Oke Ila Orangun | 7.9946°N | 4.9974°E | 867.5 | 2.04 |
| 44 | OS4 | Ikirun | 7.9227°N | 4.6347°E | 87.30 | 1.75 |
| 45 | OS5 | Ila Orangun | 8.0121°N | 4.8988°E | 80.80 | 1.85 |
| 46 | OS6 | Ilobu | 7.9021°N | 4.5315°E | 120.3 | 1.95 |
| 47 | OS7 | Iwo | 7.6292°N | 4.1872°E | 100.6 | 2.11 |
| 48 | OS8 | Ibokun | 7.8019°N | 4.7692°E | 193.8 | 2.10 |

State Code: OD→Ondo

Majority of the DNA samples extracted for use in this experiment were pure (A_260/280_ ~ 1.8) i.e. high-quality DNA extracts, with the exception of OG2 (A_260/280_ = 1.78), OG6 (A_260/280_ = 1.76) (Table 15), EK2 (A_260/280_ = 1.74), EK3 (A_260/280_ = 1.75) (Table 18), OD2 (A_260/280_ = 1.67), OD8 (A_260/280_ = 1.62) (Table 19), and OS4 (A_260/280_ = 1.75) (Table 20), with little protein and RNA contaminants found in their DNA extracts. It was observed that *Auricularia* specimen collected from Station 8 in Ogun State had the highest quantity of pelleted DNA sample with 1,548.2ng/μL of pure concentrated nucleic acid (Table 15). The least quantity of DNA extracts was obtained from *Auricularia* mushroom samples within Station 6 in Ondo State (56.5ng/μL of Nucleic acid concentration) (Table 19).

#### 3.2.2 Genetic diversity (speciation) of the sequenced *Auricularia* sp

The major allele frequency, number of alleles, genetic diversity and polymorphic information content (PIC) of the sequenced DNA extracts from all the *Auricularia* mushroom specimens geo-tagged within Southwest, Nigeria was presented in Table 21. The allele frequency ranged from 0.3542 (OPB-15) to 0.6042 (0PH-15), while the genetic diversity was from 0.5930 (0PH-15) to 0.7977 (OPB-12) and the polymorphic information content was from 0.5594 (OPH-15) to 0.7819 (OPB-12). The percentage polymorphic amplicons varied from 55.9 (OPH-15) - 78.2% (OPB-12). Therefore, OPB-12 RAPD primer gave the highest level of polymorphism (78.2%) while OPH-15 gave the least level of polymorphism (55.9%) as represented in Table 21. Nevertheless, the polymorphisms revealed by the 14 decamer primers indicate that they are good and reliable for genetic diversity assessment in Mushroom and there is a high degree of diversity in the species studied.

**Table 21:** Genetic diversity of the sequenced *Auricularia* specimens from Southwest, Nigeria

| DNA Primers | Major Allele Freq. | No. of Allele | Genetic diversity | PIC | Polymorphic Amplicons (%) |
| --- | --- | --- | --- | --- | --- |
| OPB-11 | 0.44 | 14.0 | 0.78 | 0.76 | 76.2 |
| OPB-12 | 0.40 | 13.0 | 0.80 | 0.78 | 78.2 |
| OPB-15 | 0.35 | 11.0 | 0.79 | 0.76 | 76.4 |
| OPB-20 | 0.44 | 14.0 | 0.78 | 0.76 | 76.3 |
| OPB-21 | 0.54 | 16.0 | 0.69 | 0.68 | 67.9 |
| OPH-03 | 0.46 | 12.0 | 0.75 | 0.74 | 73.6 |
| OPH-05 | 0.56 | 6.0 | 0.63 | 0.60 | 60.1 |
| OPH-10 | 0.44 | 5.0 | 0.72 | 0.68 | 67.9 |
| OPH-15 | 0.60 | 5.0 | 0.59 | 0.56 | 55.9 |
| OPT-01 | 0.46 | 11.0 | 0.74 | 0.71 | 71.3 |
| OPT-05 | 0.54 | 8.0 | 0.65 | 0.62 | 62.0 |
| OPT-07 | 0.46 | 7.0 | 0.72 | 0.69 | 68.7 |
| OPT-10 | 0.46 | 16.0 | 0.76 | 0.75 | 75.4 |
| OPT-19 | 0.52 | 14.0 | 0.70 | 0.69 | 68.7 |
| Average | 0.48 | 10.9 | 0.72 | 0.70 | 70.0 |

Sample size (n = 48)

#### 3.2.3 DNA fingerprinting and nucleotide polymorphism

The DNA marker OPB-21 had the highest number of polymorphic nucleotide (45/48) formed at 900bp (Fig 6 and Table 22), while DNA markers OPB-11 and OPB-15 at a joint highest record of polymorphic nucleotide units (46/48 each) at 100bp (Table 22). Majority of the *Auricularia* samples profiled on electrophoresis gel had no polymorphic nucleotides formed between 800-900bp units for the DNA markers OPB-11 (Fig 2), OPB-12 (Fig 3), OPB-15 (Fig 4), OPH-3, OPH-5 (Table 22), OPH-10 (Fig 7), OPH-15, OPT-1, OPT-5, OPT-7, OPT-10 (Table 22), and OPT-19 (Fig 8). A breakdown of the electrophoresis gel analysis for all base pair units was as follows: OPT-5 marker had 44/48 polymorphic nucleotides at 200bp, OPH-5 had 36/48 at 300bp, OPT-19 had 41/48 at 400bp, OPT-1 had 44/48 at 500bp, OPT-10 had 41/48 at 600bp, OPB-20 had 43/48 at 700bp and 46/48 at 800bp respectively as shown in Table 22 (only the highest number of polymorphic nucleotide was captioned).

**Fig 2.**
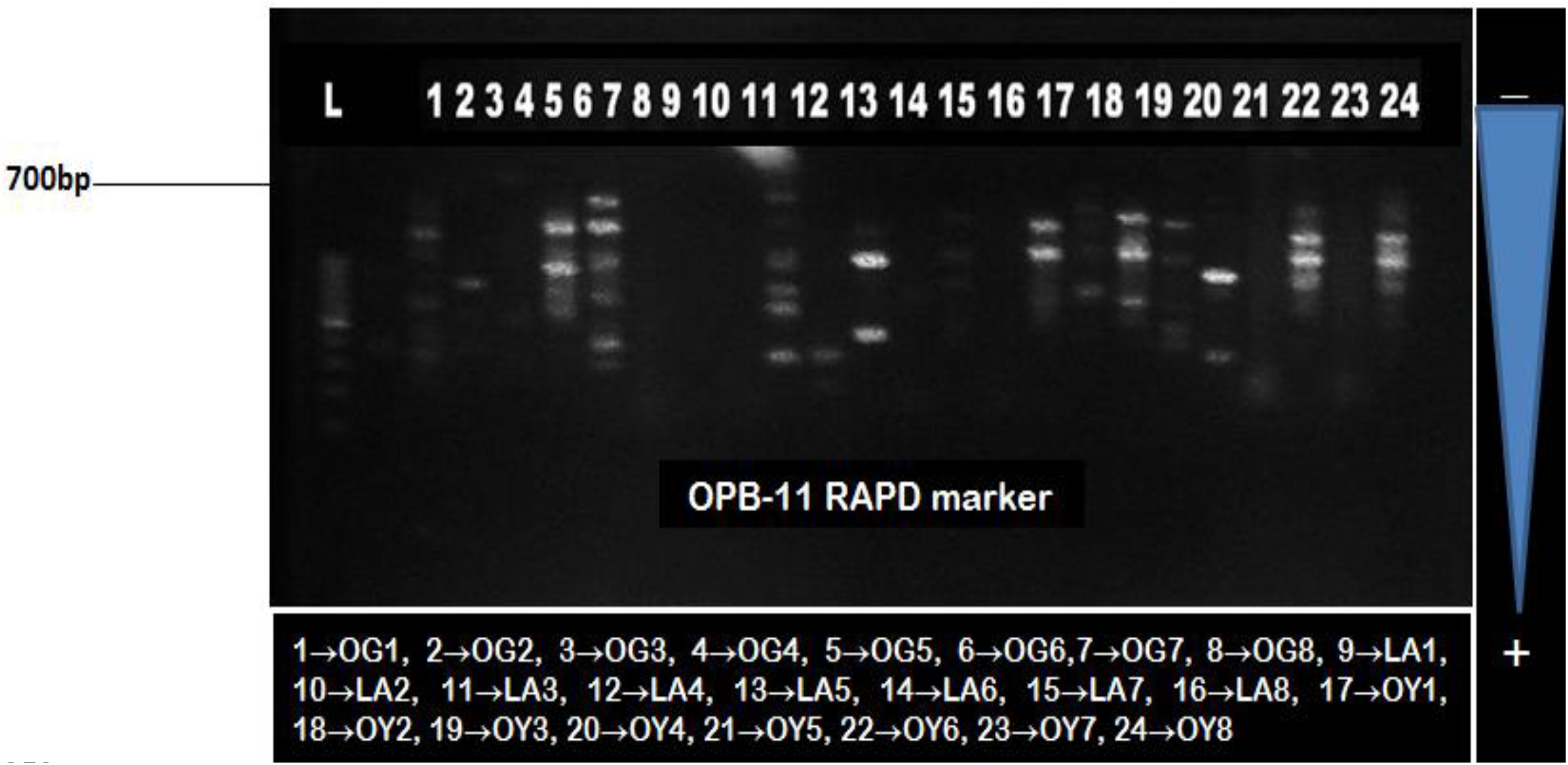
RAPD profiling of 24 *Auricularia* mushroom specimens using OPB-11 marker

**Fig 3.**
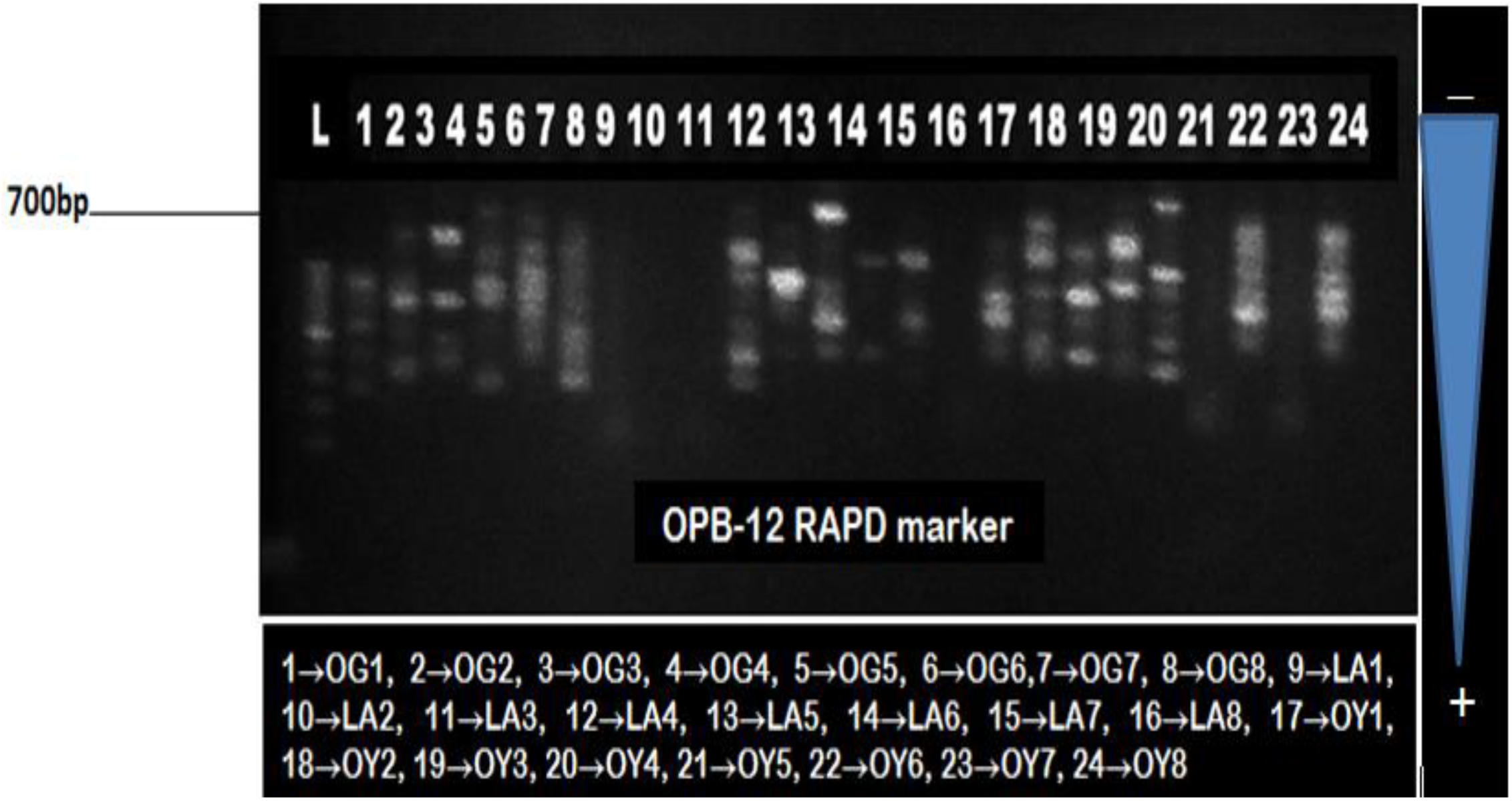
RAPD profiling of 24 *Auricularia* mushroom specimens using OPB-12 marker

**Fig 4.**
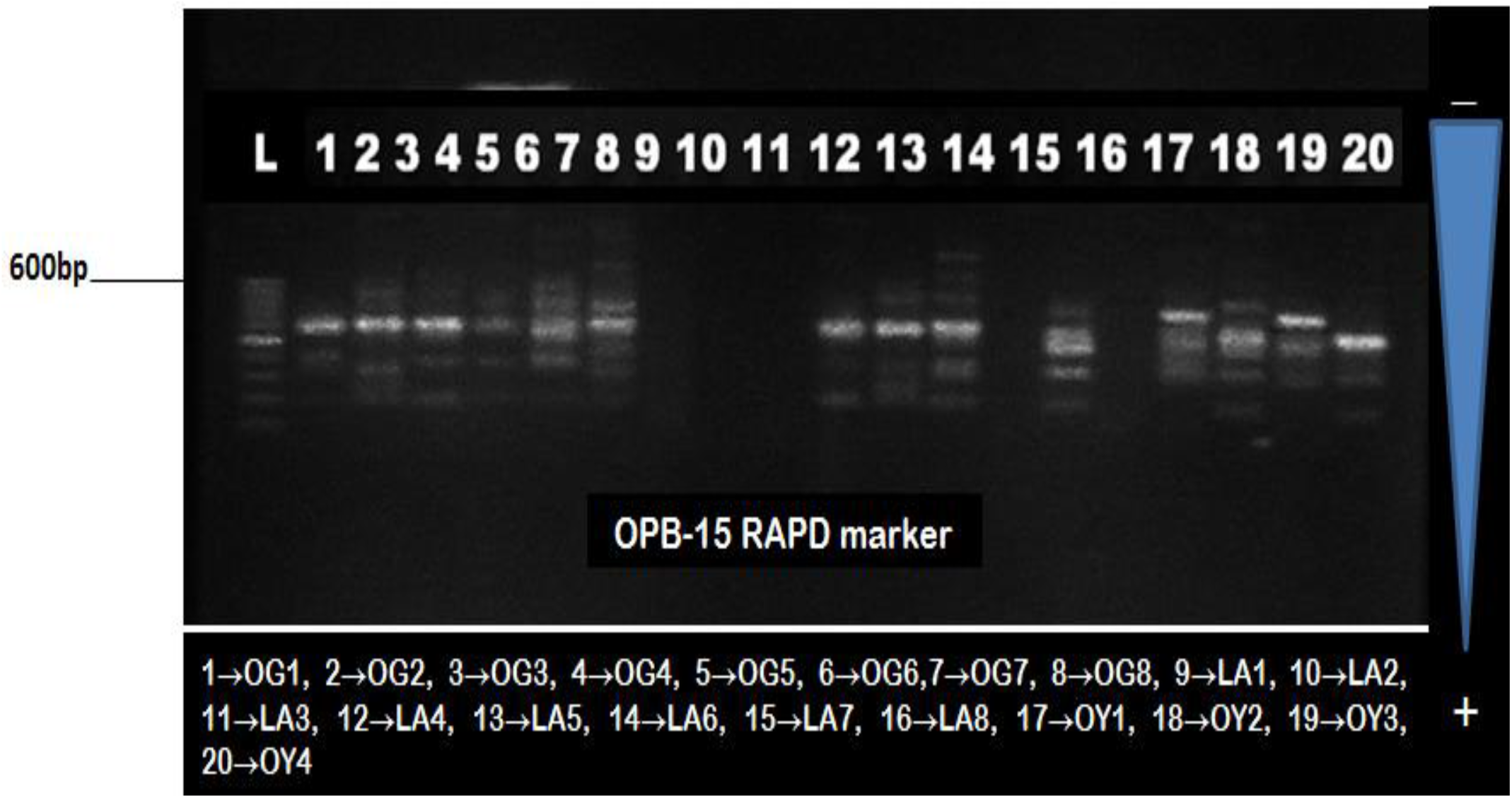
RAPD profiling of 20 *Auricularia* mushroom specimens using OPB-15 marker

**Table 22:** The number of polymorphic nucleotide amplified by different RAPD markers

| Markers | Polymorphic Nucleotide Count/bp Units |  |  |  |  |  |  |  |  |
| --- | --- | --- | --- | --- | --- | --- | --- | --- | --- |
|  | 900bp | 800bp | 700bp | 600bp | 500bp | 400bp | 300bp | 200bp | 100bp |
| OPB-11 | 00 | 00 | 06 | 13 | 13 | 06 | 33 | 38 | 46 |
| OPB-12 | 00 | 00 | 22 | 02 | 31 | 37 | 24 | 32 | 32 |
| OPB-15 | 00 | 00 | 00 | 32 | 26 | 21 | 10 | 42 | 46 |
| OPB-20 | 00 | 46 | 43 | 02 | 35 | 16 | 02 | 32 | 44 |
| OPB-21 | 45 | 09 | 12 | 37 | 17 | 09 | 01 | 09 | 39 |
| OPH-3 | 00 | 00 | 00 | 00 | 36 | 27 | 14 | 27 | 40 |
| OPH-5 | 00 | 00 | 00 | 00 | 00 | 00 | 36 | 35 | 04 |
| OPH-10 | 00 | 00 | 00 | 00 | 00 | 03 | 21 | 31 | 07 |
| OPH-15 | 00 | 00 | 00 | 00 | 00 | 03 | 04 | 07 | 05 |
| OPT-1 | 00 | 00 | 00 | 09 | 44 | 28 | 09 | 38 | 33 |
| OPT-5 | 00 | 00 | 00 | 00 | 00 | 02 | 10 | 44 | 17 |
| OPT-7 | 00 | 00 | 00 | 00 | 02 | 02 | 04 | 26 | 31 |
| OPT-10 | 00 | 00 | 38 | 41 | 15 | 12 | 05 | 04 | 26 |
| OPT-19 | 00 | 00 | 00 | 30 | 38 | 41 | 16 | 12 | 05 |

Also, it was noted that OPH-5 DNA marker expressed polymorphism in 3 bands only (100, 200 and 300bp, respectively) making it the least effective marker for this experiment, OPH-10, OPH-15 and OPT-5 all expressed nucleotide polymorphism in 4 bands each (100, 200, 300 and 400bp, respectively), whereas, OPH-3 and OPT-7 showed nucleotide polymorphism at 100, 200, 300, 400 and 500bp, respectively (i.e. 5 bands each). Furthermore, OPB-15, OPT-1 and OPT-19 expressed polymorphism in 6 band units each (100, 200, 300, 400, 500 and 600bp, respectively), OPB-11, OPB-12 and OPT-10 each expressed nucleotide polymorphism in 7 bands (100, 200, 300, 400, 500, 600 and 700bp, respectively), and OPB-20 (shown in Fig 5) had 8 band units of polymorphic nucleotides (100, 200, 300, 400, 500, 600, 700 and 800, respectively). Finally, the highest band expression for polymorphic nucleotides was found in the DNA marker OPB-21 with 9 band units (100, 200, 300, 400, 500, 600, 700, 800 and 900bp, respectively) thus, making it the most efficient marker for determining genetic variation with the earmarked *Auricularia* species in Southwest, Nigeria (Table 22).

**Fig 5.**
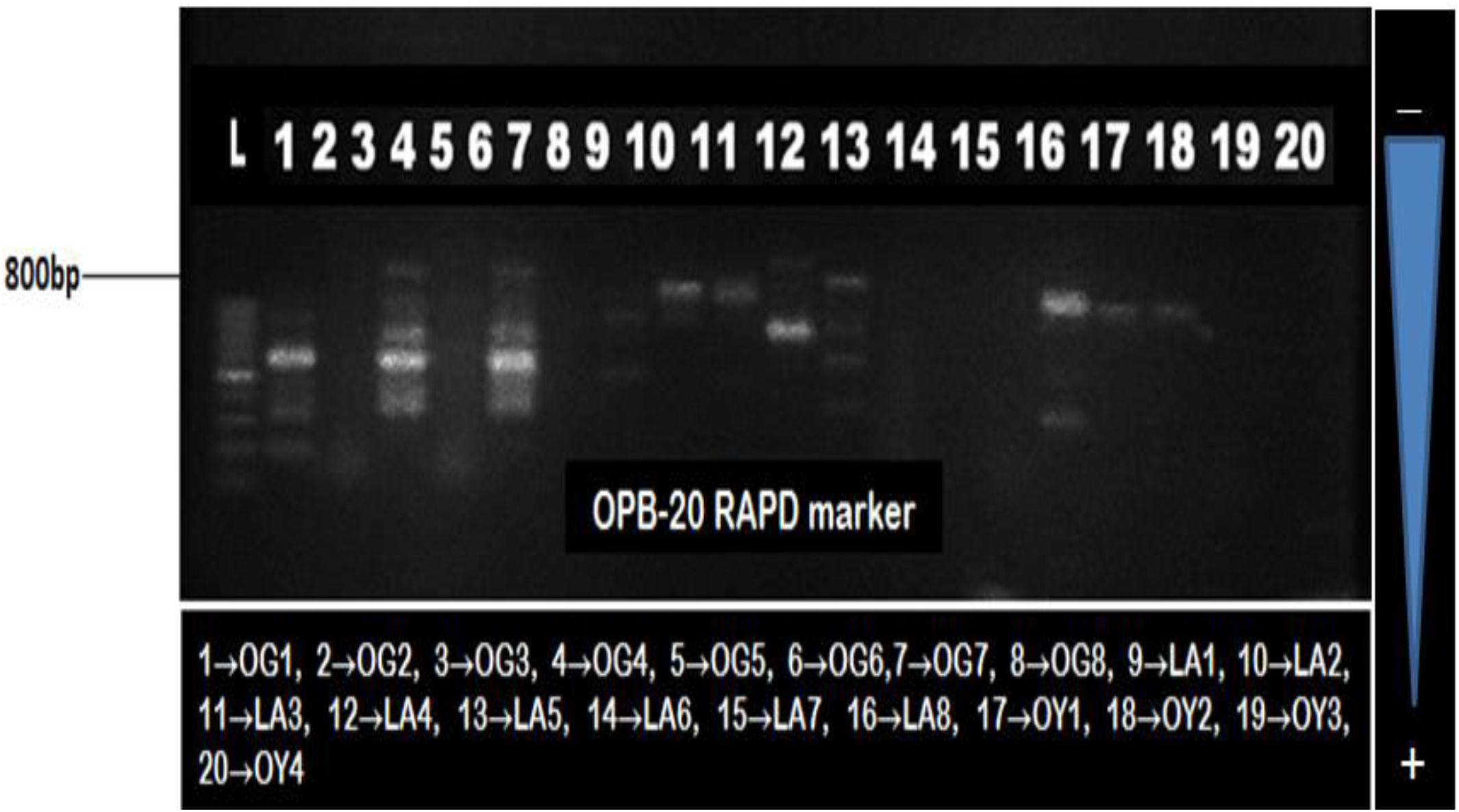
RAPD profiling of 20 *Auricularia* mushroom specimens using OPB-20 marker

**Fig 6.**
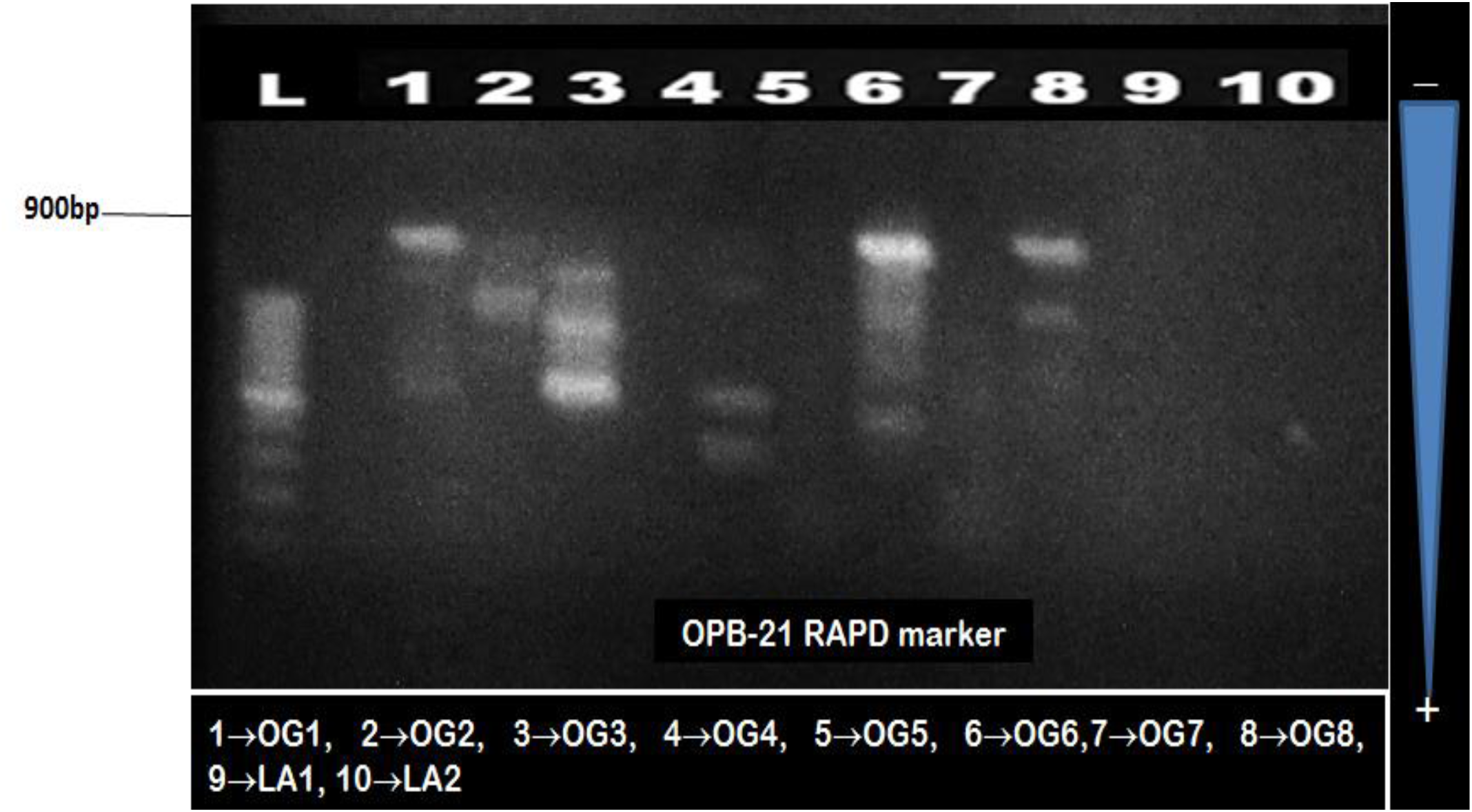
RAPD profiling of 10 *Auricularia* mushroom specimens using OPB-21 marker

**Fig 7.**
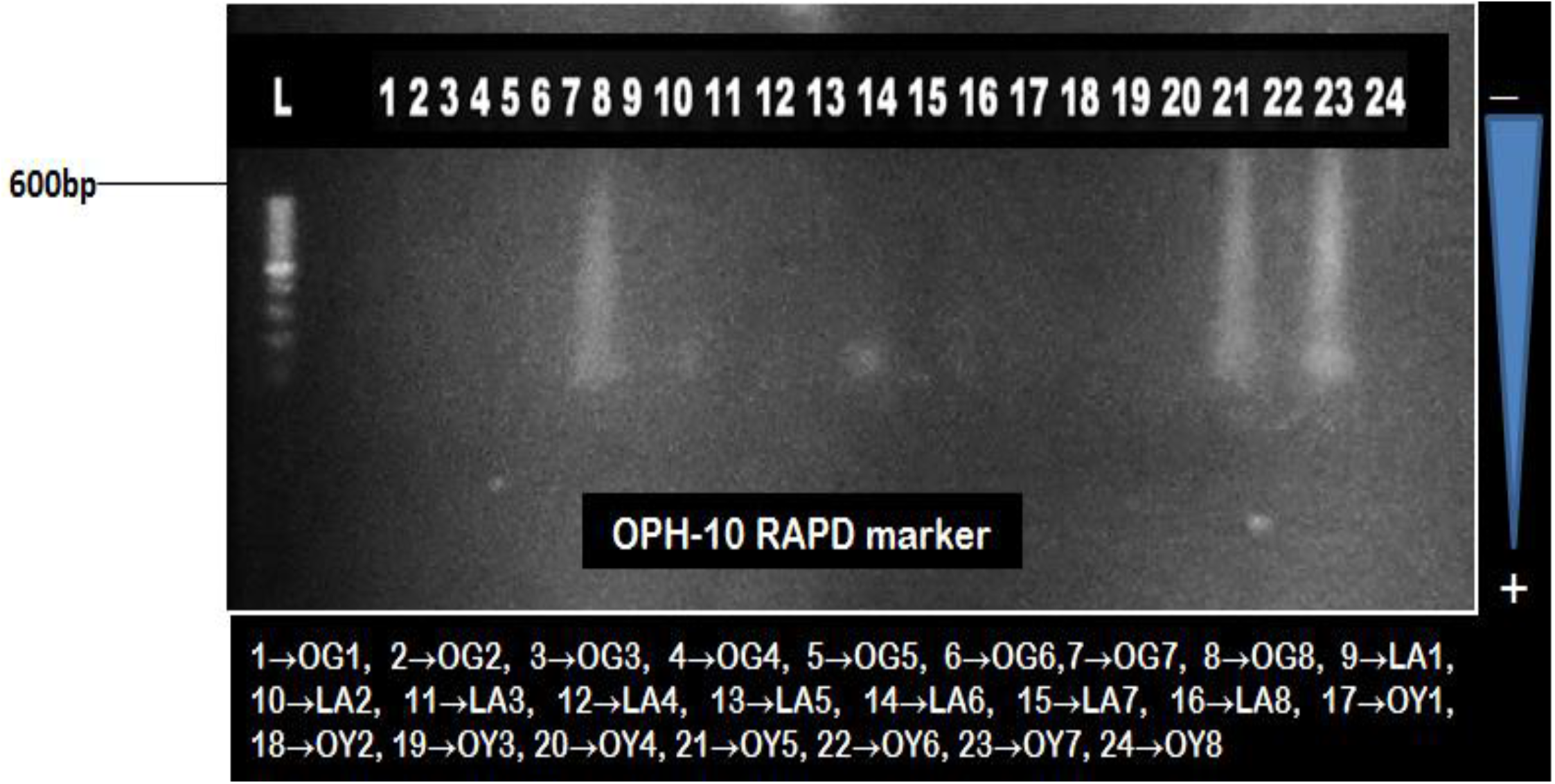
RAPD profiling of 24 *Auricularia* mushroom specimens using OPH-10 marker

**Fig 8.**
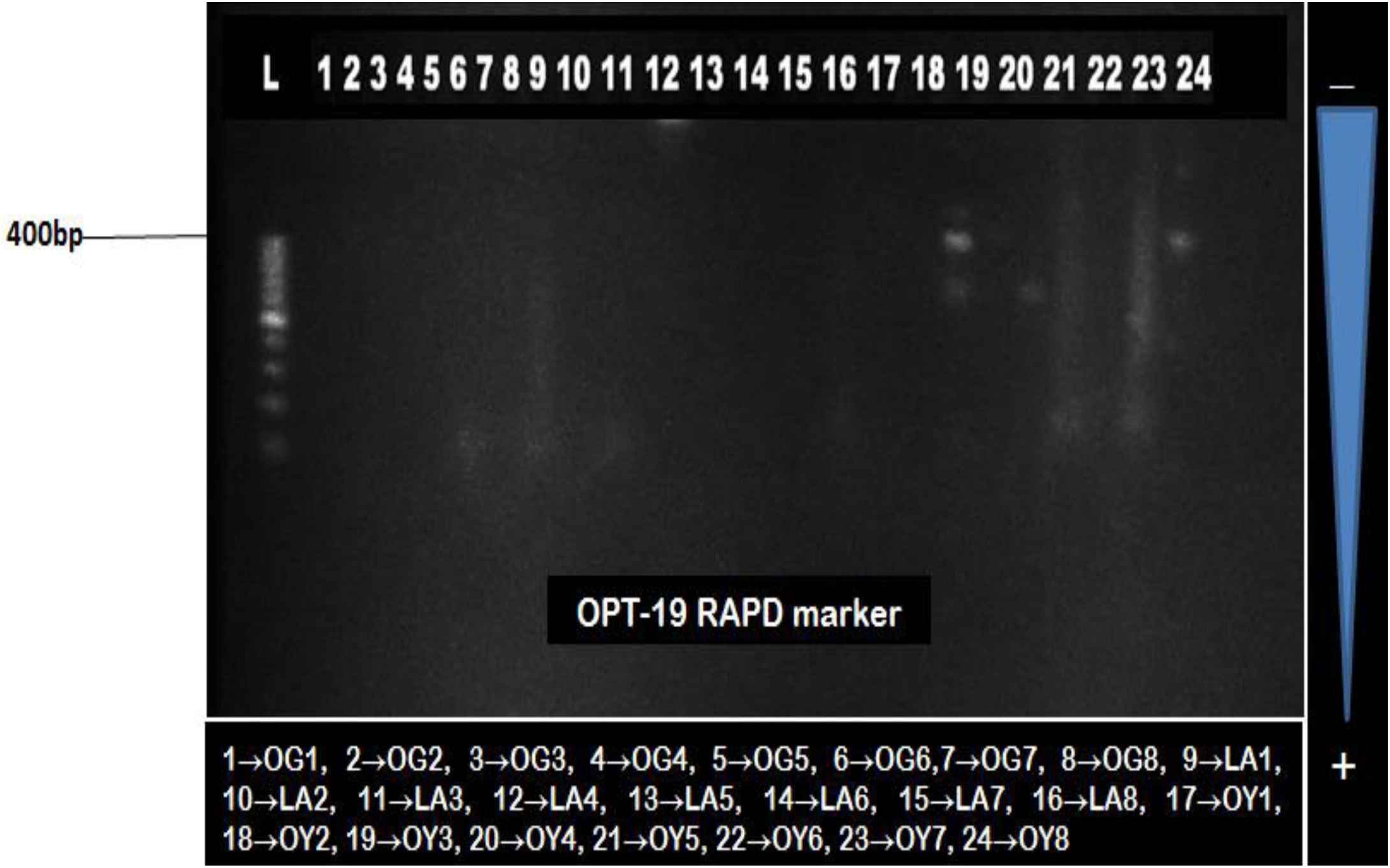
RAPD profiling of 24 *Auricularia* mushroom specimens using OPT-19 marker

#### 3.2.4 The Level of polymorphism within the *Auricularia* sp

It was observed from the records in Table 23 that the DNA markers OPB-11, OPB-15, OPB-20, OPB-21, OPT-1 and OPT-5 was able to effectively detect 90-99% polymorphism in the DNA strands of the *Auricularia* specimen profiled on electrophoresis gel (Table 23), as such they were listed as the best markers for this research. Other DNA markers such as OPH-3, OPT-10 and OPT-19 showed an impressive 80-89% variability within the *Auricularia* population in Southwest, Nigeria, while OPB-12 and OPH-5 were able to give between 70-79% variation in the examined *Auricularia* mushroom population. DNA markers OPH-10 and OPT-7 each gave between 60-69% variation, while OPH-15 was only able to detect between 10-19% variation at maximum in the Auricularia mushroom population of Southwest, Nigeria (Table 23).

**Table 23:** The percentage variation (polymorphism) that exist among the selected *Auricularia* spp

| Marker | DNA Polymorphism (%) |  |  |  |  |  |  |  |  |
| --- | --- | --- | --- | --- | --- | --- | --- | --- | --- |
|  | 900bp | 800bp | 700bp | 600bp | 500bp | 400bp | 300bp | 200bp | 100bp |
| OPB-11 | 0.0 | 0.0 | 12.5 | 27.1 | 27.1 | 12.5 | 68.8 | 79.2 | 95.8 |
| OPB-12 | 0.0 | 0.0 | 45.8 | 4.2 | 64.6 | 77.1 | 50.0 | 66.7 | 66.7 |
| OPB-15 | 0.0 | 0.0 | 0.0 | 66.7 | 54.2 | 43.8 | 20.8 | 87.5 | 95.8 |
| OPB-20 | 0.0 | 95.8 | 89.6 | 4.2 | 72.9 | 33.3 | 4.2 | 66.7 | 91.7 |
| OPB-21 | 93.8 | 18.8 | 25.0 | 77.1 | 35.4 | 18.8 | 2.1 | 18.8 | 81.3 |
| OPH-3 | 0.0 | 0.0 | 0.0 | 0.0 | 75.0 | 56.3 | 29.2 | 56.3 | 83.3 |
| OPH-5 | 0.0 | 0.0 | 0.0 | 0.0 | 0.0 | 0.0 | 75.0 | 72.9 | 8.3 |
| OPH-10 | 0.0 | 0.0 | 0.0 | 0.0 | 0.0 | 6.3 | 43.8 | 64.6 | 14.6 |
| OPH-15 | 0.0 | 0.0 | 0.0 | 0.0 | 0.0 | 6.3 | 8.3 | 14.6 | 10.4 |
| OPT-1 | 0.0 | 0.0 | 0.0 | 18.8 | 91.7 | 58.3 | 18.8 | 79.2 | 68.8 |
| OPT-5 | 0.0 | 0.0 | 0.0 | 0.0 | 0.0 | 4.2 | 20.8 | 91.7 | 35.4 |
| OPT-7 | 0.0 | 0.0 | 0.0 | 0.0 | 4.2 | 4.2 | 8.3 | 54.2 | 64.6 |
| OPT-10 | 0.0 | 0.0 | 79.2 | 85.4 | 31.3 | 25.0 | 10.4 | 8.3 | 54.2 |
| OPT-19 | 0.0 | 0.0 | 0.0 | 62.5 | 79.2 | 85.4 | 33.3 | 25.0 | 10.4 |

#### 3.2.5 Grouping of *Auricularia* sp. in Nigeria based on genotype

The population of all the *Auricularia* mushrooms currently present in the six (6) States of Southwest, Nigeria were effectively classified into six (6) clusters on the genetic dissimilarity chart (Fig 9) using PCR and RAPD markers on representative samples collected during field survey. The six (6) clusters of mushroom categories were effectively characterized into three (3) distinct species and further sub-classified into five (5) cultivars (sub-species). The genetic relatedness of all the *Auricularia* mushrooms’ population in Southwest, Nigeria was represented in Fig 9 and as such, classified thus:

**Species 1:** ***Auricularia polytricha***
  → Cultivar I (Group I): OD1, OD8, OY1, OG1, OG2, LA6, LA7, LA8
  → Cultivar II (Group II): OD2, OD3, OD4, OD5, OD6, OD7
  → Cultivar III (Group IV): LA5

**Species 2:** ***Auricularia auricula***
  → Cultivar I (Group III): OG3, OG4, OG5, OG6, OG7, OG8, OS1, OS2, OS3, OS4, OS5, OS6, OS7, OS8, EK1, EK2, EK3, EK4, EK5, EK6, EK7, EK8
  → Cultivar II (Group V): OY2, LA1, LA2, LA3, LA4, OY5, OY6, OY7, OY8

**Species 3: Unrelated *Auricularia* specimen (Outliers)**
  OY3, OY4

**Note***: The farther apart the Auricularia mushrooms on the genetic dissimilarity tree, the more related the species*.

**Fig 9.**
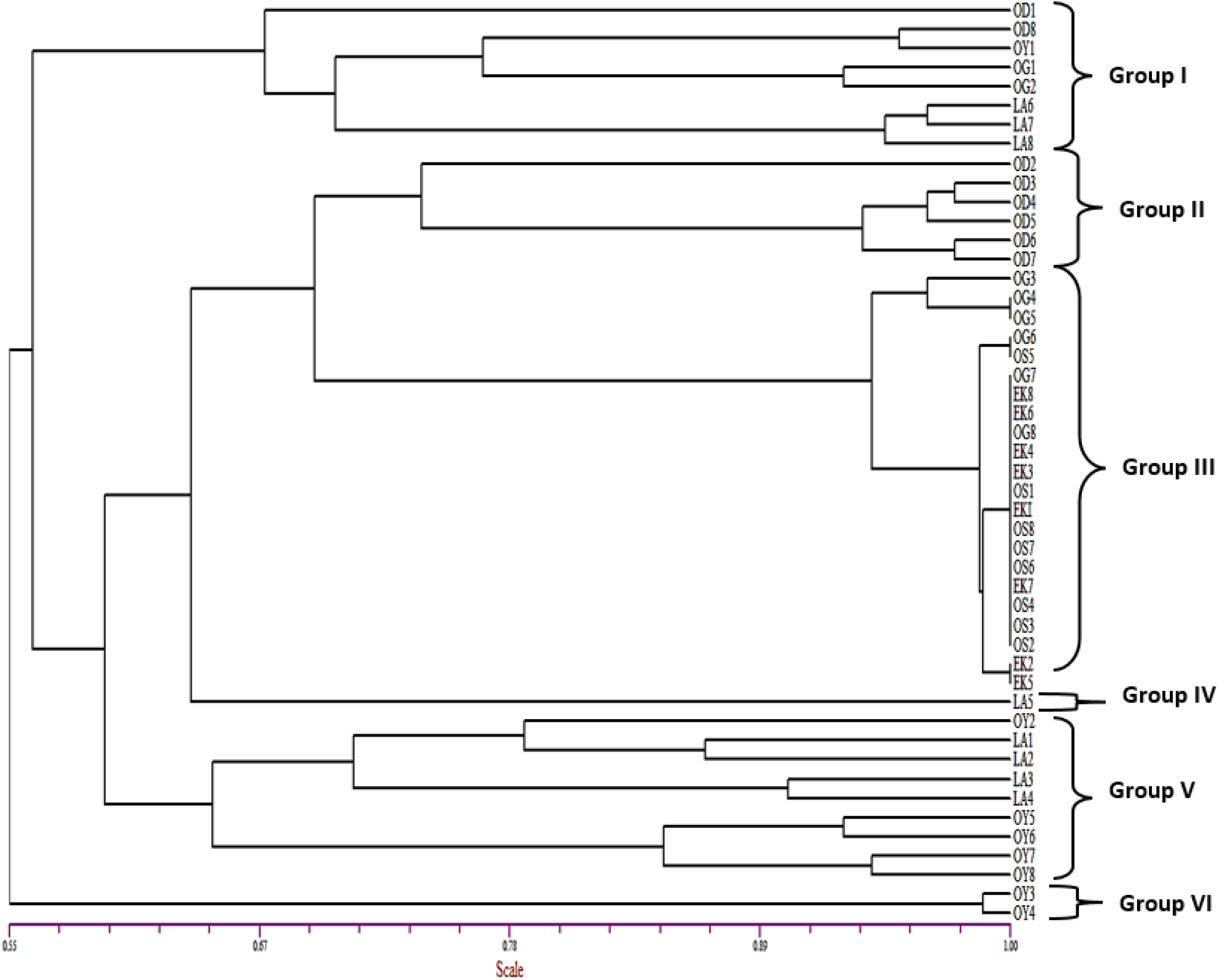
Genetic dissimilarity among the population of *Auricularia* spp in Southwestern Nigeria

## 4.0 Discussion

The morphological markers used in this study was able to identify thirty one (31) locations in Southwest, Nigeria where *Auricularia auricula* can be found, and twelve (12) locations where *Auricularia polytricha* thrived better within the region under survey. It was noted also that six (6) out of the earmarked fifty four (54) locations had no *Auricularia* mushrooms present within their domain. These could be as a result of severe foraging (wild mushroom exploit) or destruction of their natural habitats by man. Samples of *Auricularia auricula* was evenly distributed in Ekiti, Osun, Ogun, Oyo and Lagos States; but there was none identified in Ondo State as at the time of filing this reports. There are limited scientific explanations to this observation since the region enjoys a seemingly even distribution of rainfall and sunshine as do other States in Southwestern Nigeria. *Auricularia polytricha* was found in abundance in Ondo and Lagos States only, but none was found in Ogun, Ekiti, Osun and Oyo States. Ironically, a scientific explanation is imminent. Therefore, more research work is recommended in this field and with regards to the observations outlined by this research in order to fully address the questions raised. The findings were in line with the reports of Onyango *et al*., 2010) who identified three (3) main strains (brown, dark brown and yellow brown) of *Auricularia* mushrooms occurring in the forest region of Africa using morphological markers. Also, Li *et al*. (2011) reported that similar clustered patterns, reveals that all the tested strains could be divided into three distinct groups, each of which was correlated with different geographical regions.

In order to ascertain and fully establish the genomic differences that exist among the mushroom specimens based on the influence of the environment and geographical boundaries, and further enhance the characterization made in this research based on morphological markers. The mushrooms were further subjected to molecular testing using PCR and RAPD techniques. Molecular markers such as rDNA sequencing, Restriction fragment length polymorphism (RFLP), Random amplified polymorphic DNA (RAPD) and genotyping have been used to discriminate mushroom species or strains of *Agaricus, Auricularia, Ganoderma, Lentinula, Stropharia*, and *Volvariella*. All of these technologies provided data for mushroom strain identification and protection (Chandra *et al*., 2010).

The DNA marker “OPH-5” was the least effective marker for this experiment, while OPB-21 was the most efficient marker for determining genetic variation with the earmarked *Auricularia* species in Southwest, Nigeria. This was in agreement with the research of Khan *et al*., (2011) who conducted molecular characterization of Oyster mushroom (*Pleurotus* spp.) using 14 RAPD primers and obtained the highest polymorphism by primers OPL-3 (72.70 %) and OPL-11 (70%). Two species (P-56 and P-17) were found to be genetically similar having a similarity value of 86%. The population of all the *Auricularia* mushrooms currently present in the six (6) States of Southwest, Nigeria were effectively classified into six (6) clusters on the genetic dissimilarity chart using PCR and RAPD markers on representative samples collected during field survey. The six (6) clusters of mushroom categories were effectively characterized into three (3) distinct species and further sub-classified into five (5) cultivars (sub-species). The result obtained in this study also agrees with the report of Ravash *et al*., (2009) who used RAPD markers to confirm the similarity or dissimilarity of genetic relationship of *Pleurotus* spp.

## Conclusion

The use of morphological markers only for characterization of *Auricularia* species found in Southwest, Nigeria was pragmatic as it produced a very good result but the best option was a combination of both morphological and molecular markers (PCR and RAPD) to determine the genetic diversity and variation within the large genomic entity of *Auricularia* mushroom population in Southwest, Nigeria.

## Ethical Statement

This is to confirm that:

Prof. Clementina O. Adenipekun, Dr. V. S. Ekun, Dr. P. M. Etaware and Dr. Omena B. Ojuederie declares that they have no conflict of interest and that they actively participated in the research both in the field and in the procurement of materials for morphological and molecular analysis.

Thank you

Peter M. Etaware (Ph.D.)

## Ethical Approval

‘Not applicable’.

## Consent to Participate

All the authors gave their consent to participate in this research.

## Consent for Publication

All the authors unanimously agreed that this article should be published

## Availability of Data and Material

All data and material are present in this publication.

## Authors’ Contributions

E.V.S. and A.C.O conceptualized and designed the experiment. E.V.S conducted the research and E. P. M. wrote the draft manuscript. E. P. M. and O.O.B. reviewed the manuscript. All authors approved the final version of the manuscript.

